# Positive effects of larval quality on early barnacle recruitment through adaptive substrate selection

**DOI:** 10.64898/2026.10.19.760179

**Authors:** PGM Guerra, ACC Barbosa, R Tremblay, AAV Flores

## Abstract

Variation in larval supply alone often fails to explain recruitment dynamics in marine benthic populations, suggesting that larval traits may influence settlement success. We tested whether larval quality influences habitat selection and recruitment in the intertidal barnacle *Chthamalus bisinuatus*. Over a 26-day field sampling, we found no correlation between larval supply and settlement rates on experimental arenas featuring varying microtopographies. Instead, recruitment dynamics tracked changes in larval quality. Early settlement was primarily driven by low-quality larvae, consistent with the ‘desperate larva hypothesis’, whereas later recruitment was characterized by high-quality larvae. Cyprids generally preferred rugose over smooth surfaces, but substrate selectivity depended on larval condition and disappeared after biofilm development. Early benthic performance indicated adaptive habitat selection, with higher survival and larger recruits on rugose surfaces. These findings identify larval quality as a key driver of settlement, shaping recruitment patterns and should be incorporated into ecological models of marine population connectivity.

## Introduction

Population fluctuations often result from the interplay between abiotic tolerance, species interactions, and propagule supply dynamics (Holling 1973; Eckert 2003; Moore and Stanton 2014). At a geographical scale, population dynamics are shaped by barriers to dispersal (Gaylord et al. 2000; Turnbull et al. 2000), biotic processes (Moeller et al. 2012), and environmental quality (Eckhart et al. 2011). Species exhibiting patchy distributions in apparently continuous habitats provide excellent model systems for studying spatial ecology, as they allow the investigation of dispersal, habitat quality, and biotic interactions interact to determine spatial occupancy (Hastings 1991; Doebeli 1995; Moore 2009; Sexton et al. 2009).

Theoretical models propose that dispersal between habitat patches stabilizes population sizes by reducing density fluctuations through the spread of offspring across heterogeneous habitats (Den Boer 1968; Hastings 1991; Doebeli 1995). Following this paradigm, species with prolonged dispersal phases are expected to exhibit greater adult population volatility compared to those with shorter dispersal periods (Thorson 1950). However, empirical evidence challenges this view. A comprehensive analysis of 570 time series of intertidal and subtidal benthic species by Eckert (2003) failed to consistently support these theoretical predictions. Instead, this research indicates that local environmental conditions and life-history characteristics frequently outweigh the influence of pelagic larval duration in determining population fluctuations.

The dynamics of habitat-forming species, such as corals and barnacles, are predominantly influenced by the monopolization of space and the availability of vacant habitats, often generated by episodic disturbance events (Connell 1978; Sousa 1984; Hughes et al. 2007). These limited recruitment opportunities foster intense competition for suitable settlement sites, particularly at microscale levels, where small-scale variation in substrate features significantly modulates larval settlement preferences and subsequent community structure (Raimondi 1988; Vermeij and Sandin 2008; Toupoint et al. 2012b). These processes emphasize the importance of understanding settlement processes across spatial hierarchies, ranging from broader habitat patches to the specific microhabitats ultimately colonized by larvae

Although larval supply is widely recognized as a key determinant of recruitment in marine benthic populations (Underwood and Fairweather 1989), it is often weakly correlated with early benthic recruitment (Olivier et al. 2000; Shanks 2009; Toupoint et al. 2012a). This asynchrony is attributed to larval behavior (Pineda et al. 2010), which is closely linked to lipid reserves accumulated during the pelagic phase (Thiyagarajan et al. 2002; Tremblay et al. 2007). The transition from the pelagic phase to benthic attachment involves active microhabitat selection (Miron et al. 2000), with larvae relying on these reserves to sustain swimming, habitat selection and metamorphosis (Pechenik et al. 1998; Jarrett 2003; Thiyagarajan et al. 2002). Poor-quality larvae, characterized by reduced reserves, may fail to identify or colonize favorable microhabitats, whereas high-quality larvae are better equipped to explore and establish in these key settlement patches (Lucas et al. 1979;Pechenik et al. 1998; Jarrett 2003). This contrast is central to the ’desperate larva hypothesis’ (Knight-Jones 1953), which predicts that energy-depleted larvae progressively lose their ability to select optimal settlement habitats and instead settle opportunistically on whichever substrate they encounter, potentially compromising post-settlement survival and growth (Rittschof et al. 1984; Toonen and Pawlik 1994; Tremblay et al. 2007).

Given the scarcity of suitable recruitment sites, particularly for species with strong space-monopolization traits, larval quality is a critical determinant of microhabitat selection and subsequent recruitment success (Pechenik et al. 1998; Jarrett 2003; Thiyagarajan et al. 2002). Because even small-scale substrate variations significantly modulate settlement, survival and growth (Raimondi 1988; Vermeij and Sandin 2008; Toupoint et al. 2012b), understanding the interplay between larval quality, microhabitat availability and settlement success provides critical insights into the population dynamics of habitat-forming species (Pechenik et al. 1998). Here, we test the hypothesis that variability in larval condition, measured by lipid content, dictates the ability of larvae to select and settle in favorable microhabitats. Using a field experiment designed to assess barnacle larval habitat selectivity across contrasting microtopographies, we explore the implications of larval quality for recruitment outcomes and early benthic performance.

## Materials and Methods

### Study area

All field work was conducted along a rocky coastline within 5 km from the Centre of Marine Biology of the University of São Paulo (Fig. 1). The area is situated in a broad transitional zone along the Southwestern Atlantic (Appendix S1), where temperate and tropical species coexist (e.g., Martins et al. 2007; Benkendorfer and Soares-Gomes 2009). The study coastline faces the São Sebastião Channel (SSC, Fig. 1B) and is therefore relatively sheltered, occasionally exposed to moderate swell due to southwesterly winds. Such remote forcing, along with the intrusion of South Atlantic Central Waters during the summer (Ciotti et al. 2010), may promote vertical mixing, fertilizing surface waters otherwise oligotrophic (Aidar et al. 1993).

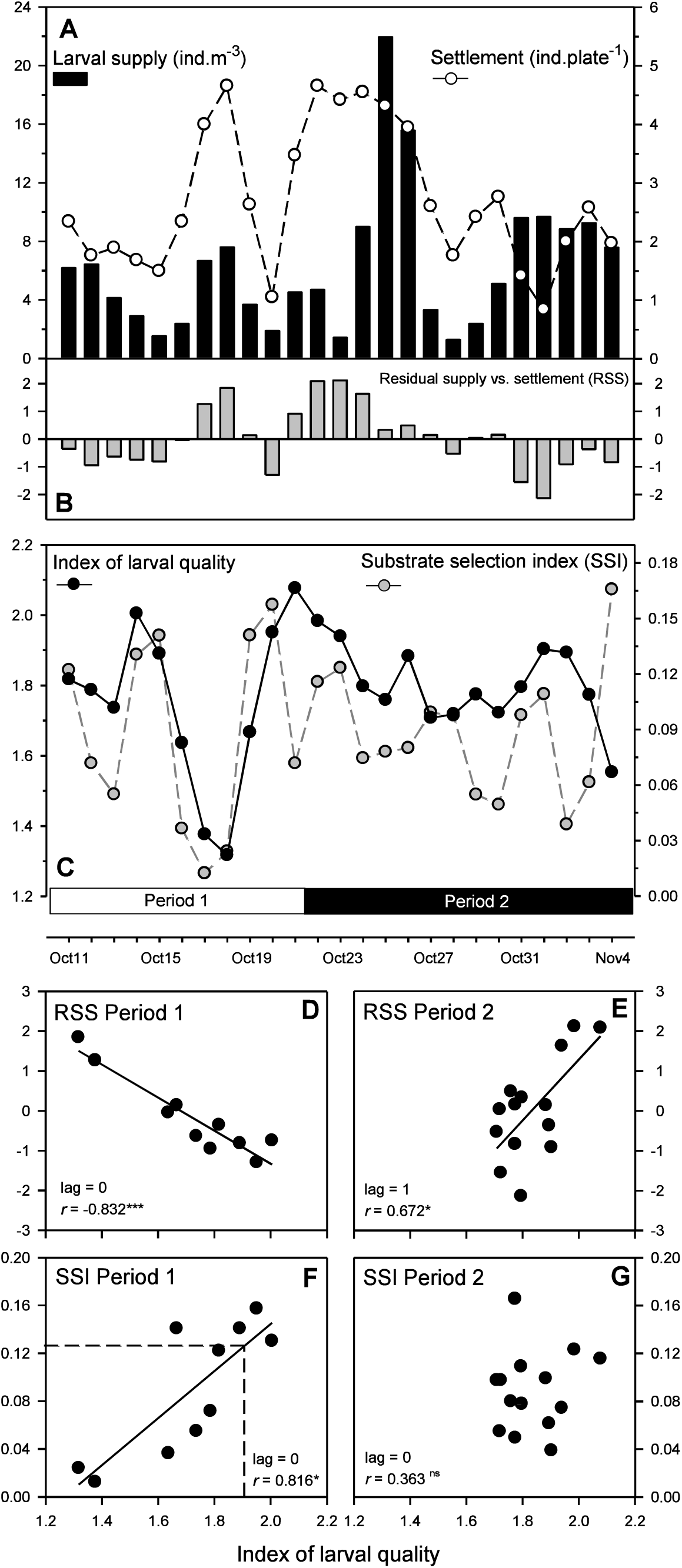

### Biological model

*Chthamalus bisinuatus* (Pilsbry 1916) is the most abundant intertidal barnacle species in subtropical rocky shores along the southwestern Atlantic coast (Young 1993). Under moderate wave exposure, this species forms dense populations that often saturate available space at the mid-shore level (80-100% substrate cover; Bueno et al. 2010; Kasten and Flores 2013). As in other thoracic barnacles, larval development comprises six naupliar planktotrophic stages and a non-feeding cypris stage. This latter stage relies exclusively on energetic reserves to maintain swimming, habitat selection and perform metamorphosis.

### Larval supply and quality, benthic microhabitat selection and post-settlement performance

#### Nearshore plankton sampling

A submersible water pump, installed at an average 1.5 m depth, was used to sample the nearshore water column and provide estimates of larval supply (ind.m^-3^) and larval quality at Segredo Beach (23°49’42” S; 45°25’21” W), from October 10 to November 7, 2014. The pump operated daily from 09:00 to 11:00 and 14:00 to 16:00, filtering approximately 28.8 m^3^ of water per day. Plankton was retained in a collector mounted on a 150 µm plankton net. Cyprids were rapidly sorted from samples, counted, and assessed for quality based on lipid storage as indicator. Relative lipid abundance (RDA) was determined via image analysis (% cover of cyprid lateral area), which were highly significantly correlated to the quantity of stored triacylglycerols (TAGs; see Appendix S2). To quantify lipid content in a large number of larvae, a technician was trained for rapid discrimination of larvae among 9 different RDA categories, ranging from 1 (0-5%) to 9 (40-45%). When larval density was too high, subsamples were obtained using a Folsom plankton splitter.

### Benthic sampling

Over the same period and at the chthamalid barnacle zone just above the subtidal water pump, daily settlement rate was estimated uisng 10 x 10 cm plates. These polymethyl methacrylate (PMMA) plates were divided into four 5 x 5 cm sectors. Three of them received different microtopographies by pressing Nitex® nylon meshes of different sizes (150, 300 or 500 µm) at 160°C for 4 h, and one left unaltered (smooth). Rugose plates ended up with pit sizes of ca. 250, 430 and 680 µm, approximately matching the topographies obtained by Berntsson and coworkers (2000). These sizes were respectively, smaller, close to and larger than the average *C. bisinuatus* cyprid length. After being held under running seawater for 5 d to remove any chemical substances that could inhibit larval settlement, 19 arenas were installed on the shore, maintaining a distance of at least 30-50 cm between neighboring plates. All plates were photographed daily using a super macro lens (waterproof Nikon Coolpix AW100, 16 megapixels) to estimate settlement rates. Images were also used to track randomly selected settlers (n = 21) at preferred and non-preferred surfaces (see results) over two 5 d periods, both during major settlement events (Oct. 17-21 and Oct. 23-27) to estimate mortality and growth rates. Image analyses were performed in ImageJ®.

### Numerical and statistical analyses

Correlational evidence was tested for relationships between (i) larval supply and settlement (benthic-pelagic link); (ii) cyprid lipid content and supply-settlement residuals (testing settlement competency vs. the ‘desperate larva hypothesis’ (Knight-Jones 1953); and (iii) cyprid lipid content and substrate selectivity (testing reserve limitations on selection). Before analysis, time series data were smoothed using 2 d moving averages. We assessed significance using conventional correlation coefficients, correcting degrees of freedom to account for autocorrelation over the maximum testable lag interval (*N*/5, in which *N* is the total series length). This alternative procedure may increase test-power in cases when low-frequency variability is important (Pyper and Peterman 1998).

The full settlement dataset (n = 1,330) was used to assess settlers distribution across microtopographies using qui-square procedures, followed by three planned contrasts: (i) smooth vs. rugose (testing non-specific preference for rugose surfaces), (ii) 300/500 µm vs. 150 µm (testing preference of topographies that generate shelter); and (iii) 300 µm vs. 150/500 µm (testing selection fit-to-larval-size preference). Type I errors were Bonferroni corrected (Bland and Altman 1995). The frequency of settlers in overall preferred and non-preferred surfaces was then used to calculate daily qui-square values, setting up a time series of substrate selectivity.

To test for adaptive substrate selection, we tracked individuals tracked on experimental plates over the specified settlement events to assess early benthic performance (5 d) on preferred and non-preferred surfaces. Final size (opercular diameter, in µm) was evaluated using a 2-way mixed model ANOVA, in which ‘substrate’ (preferred, non-preferred) and ‘settlement event’ (Oct. 17-21, Oct. 23-27) were fixed and random factors, respectively. Survival (%) was analyzed using log-linear models applied to three-way contingency tables (‘substrate type’ x ‘survival’ x ‘settlement event’) (Williams 1976; Sokal and Rohlf 1995).

## Results

### Larval supply, settlement rate and larval quality

Overall, propagule density (larval supply) was a poor predictor of settlement rates at adjacent benthic grounds (Fig. 1A, Table 1). Although some correspondence occurred the mid-October supply event, and to a lesser extent between October 24-26 and the major settlement event, correlations were not significant at any of the tested lags (-5 to 5 d). The maximum correlation coefficient for the entire series was found at lag 0, but remained below the statistical threshold (Table 1). Supply pulses were more acute than settlement events, suggesting that a fraction of cyprids did not settle immediately, but accumulated in the nearshore pelagic zone. A substantial fraction of such pelagic-benthic decoupling was explained by larval quality (Table 1). Residuals of supply vs. settlement relationships (RSS; Fig. 1B) clustered in 3-6 d sets (in either direction), mirroring the temporal variation in larval quality (Fig. 1C). However, the influence of larval quality on settlement rates differed between Period 1 (October 11-21) and Period 2 (October 22 – November 4) (Fig. 1B, 1C). During Period 1, residual settlement and larval quality oscillated in opposite directions, resulting in a clear negative correlation (Fig. 1D) at lag 0; specifically, 69% of the positive residual settlement was explained by the immediate settlement of larvae with low energetic lipid content. Conversely, in Period 2, larval quality had a significant positive effect on settlement, with increased larval quality linked to higher residual settlement after one day (Fig. 1E). Thus, higher settlement rates in Period 1 were associated with larvae having low lipid reserves, whereas in Period 2, they were associated with larvae having high lipid reserves.

**Table 1.** Main results of cross-correlation analyses testing relations between (1) larval supply and larval settlement, (2) cyprid lipid contents (a proxy of larval quality) and unexplained variation of supply vs. settlement relationships, and (3) larval quality and substrate selection. In all cases, statistical results are restricted for the lag (in days) in which the highest correlation coefficient (*r*) was observed. Corrections for hypothesis testing owing to autocorrelation of time series are detailed in the main text. Statistic details for significant cases (*p_adj_* < 0.05) are highlighted in bold.

|  | Larval supply vs. Larval settlement |  |  | Larval quality vs. Res.<sub>sup</sub> vs. set |  |  | Larval quality vs. Substrate selection |  |  |
| --- | --- | --- | --- | --- | --- | --- | --- | --- | --- |
| | lag | $r$ | $p_{adj}$ | lag | $r$ | $p_{adj}$ | lag | $r$ | $p_{adj}$ |
| Total | 0 | 0.323 | 0.223 | 2 | 0.344 | 0.177 | **1** | **0.574** | **0.003** |
| Period 1 | 0 | 0.71 | 0.179 | **0** | **-0.832** | **0.005** | **0** | **0.816** | **0.048** |
| Period 2 | 2 | 0.256 | 0.399 | **1** | **0.672** | **0.012** | 1 | 0.363 | 0.302 |

### Larval quality and substrate selection

Settlement data revealed clear substrate selection (Fig. 2). Regardless of texture, cyprids settled more frequently on rugose than on smooth surfaces (_χ_^2^ = 45.5, *d.f.* = 1, *p* < 0.0001). Within rugose surfaces, no significant preferences were observed between topographies providing shelter (300 µm and 500 µm; _χ_^2^ = 0.92, *d.f.* = 1, *p* = 0.821), or those matching larval size (300 µm; _χ_^2^ = 1.32, *d.f.* = 1, *p* = 0.724). Consequently, we used a substrate selection index (SSI) based only on the settlement frequencies on smooth vs. rugose surfaces to calculate daily deviations from random distribution:

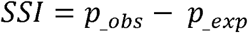

where *p*___*_obs_* represents the observed proportion of settlers on rugose surfaces, and *p*___*_exp_* denotes the expected proportion for a random distribution (= 0.750). Accordingly, SSI < 0 indicates selection for smooth surfaces (an unlikely case), SSI = 0 no selection, and SSI > 0 indicates selection for rugose surfaces. Within the latter, an SSI of 0.125 corresponds to 50% of the larvae selecting for rugose surfaces, while an SSI of 0.250 corresponds to 100% selectivity.

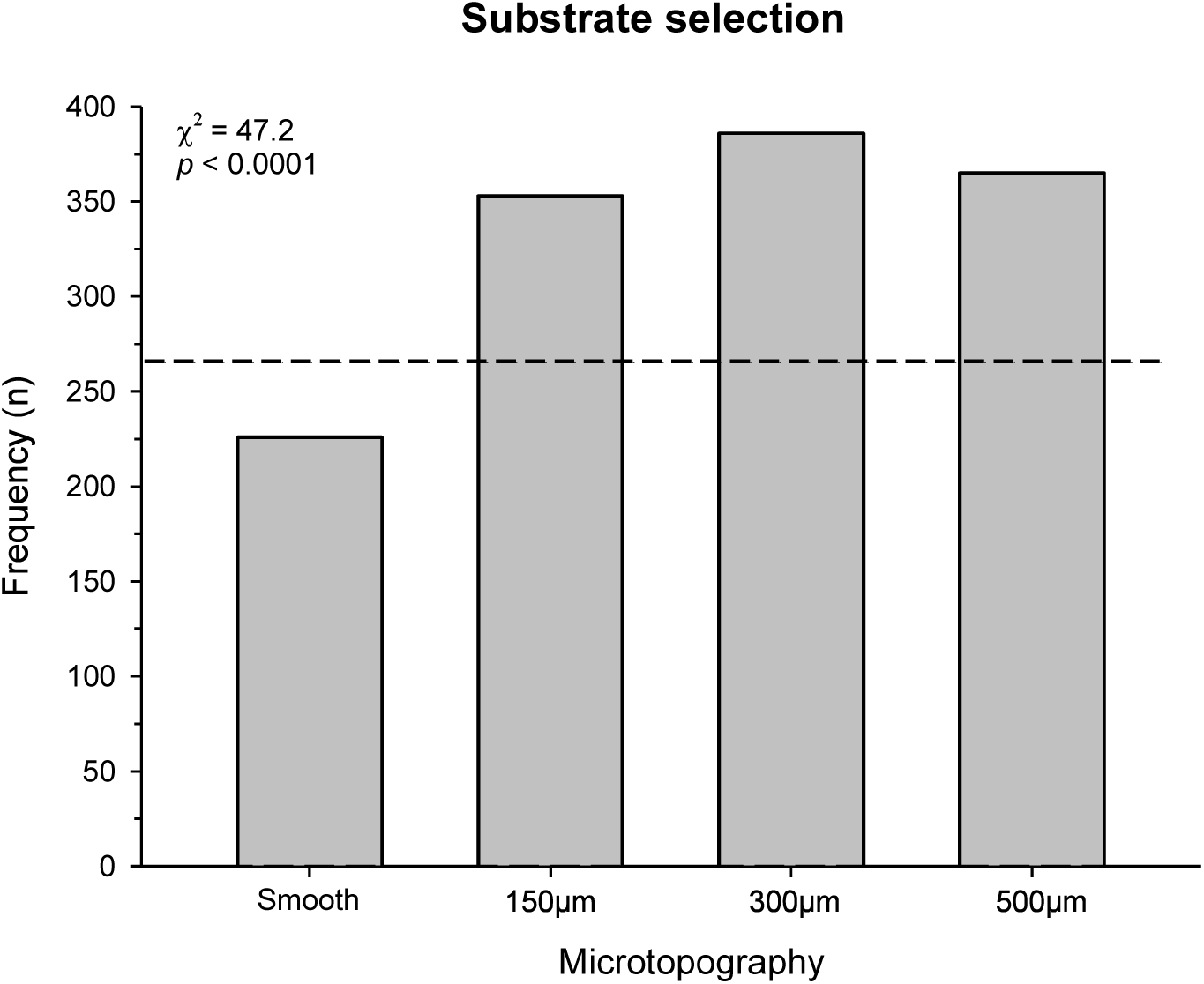

This time series is shown in Figure 1C. SSI values remained positive throughout the study, indicating a persistent preference for rugose surfaces. The average SSI was 0.088 (range: 0.012 -0.166 ), reflecting an average of 35% selective larvae (SSI_35_), varying from 5% to 66%. SSI variation was strongly and positively related to larval quality during Period 1, but not during Period 2 (Fig. 1C, F, G). During Period 1, when plates were still free from advanced biofilm coating, larvae with higher lipid contents were more selective, with larval quality explaining 67% of the SSI temporal variation (Fig. 1F). A lipid content index of 1.90 was estimated as the threshold where 50% of larvae select rugose surfaces (SSI_50_). This correlation did not persist during Period 2 (Fig. 1G).

### Adaptive value of substrate selection

Early benthic performance was consistently superior on rugose surfaces than on smooth ones across both settlement events (Fig. 3). Survival was significantly higher on rugose surfaces (70%) compared to smooth ones (32%; _χ_^2^_subxsur_ = 41.39, *p* < 0.001), with no significant interaction between substrate type and settlement event (_χ_^2^_subxsurxev_ = 0.36, *p* = 0.546). Similarly, final size was nearly 40% greater on preferred rugose substrates (c.a. 350 μm) than on smooth surfaces (ca. 250 μm; *F*_sub_ = 23.47, *p* < 0.0001), with no significant interaction between substrate and settlement event (*F*_subxev_ = 0.68, *p* = 0.413).

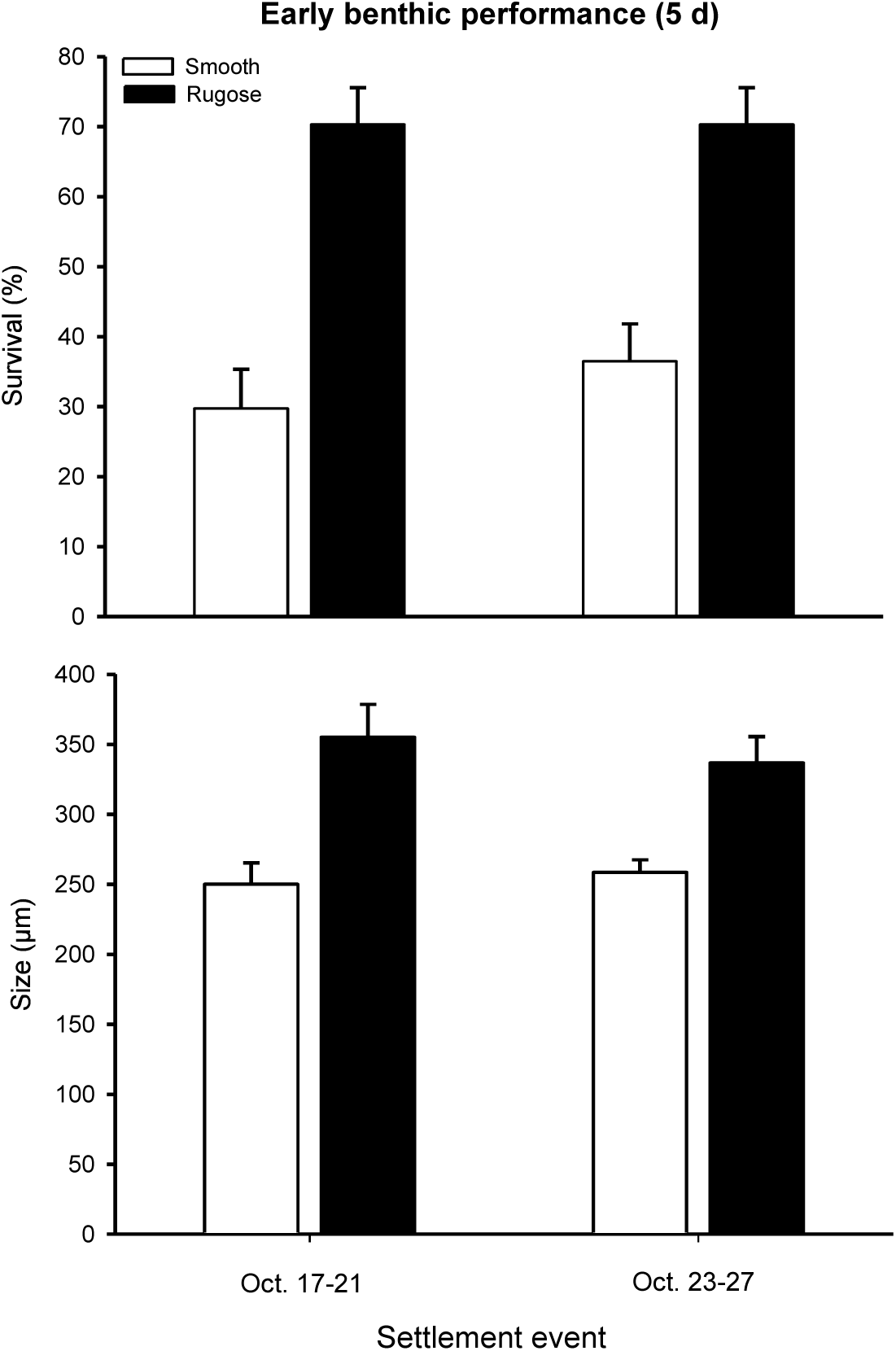

## Discussion

Our results demonstrate that larval quality is a primary determinant of habitat selection, with profound implications for subsequent benthic performance. However, two distinct temporal recruitment patterns, mediated by larval physiological condition, were observed. During the first 10 days of the study, settlement was largely explained by low-quality larvae exhibiting minimal substrate selectivity. During this period, larvae with a quality index <1.7 were frequent, a finding aligned with the ‘desperate larva hypothesis’, which posits that cyprids lose their settlement selectivity as energetic reserves are depleted (Rittschof et al. 1984; Toonen and Pawlik 1994; Tremblay et al. 2007). However, within this same period, larvae with high lipid reserves (index >1.9) displayed a clear preference for rugose surfaces and enhanced early survival and growth. This contrast illustrates that the capacity to discriminate between microhabitats, and the associated fitness benefits, is not a fixed trait of the species, but is instead physiologically dependent on the individual larval energy budget.

Habitat selection is a critical life-history bottleneck, as early post-settlement mortality for marine invertebrates often exceeds 60-99% within the first few weeks (Thorson 1950; Hunt and Scheibling 1997). Settlement by high-quality cyprids is well-documented to improve metamorphosis success and juvenile growth performance (Jarrett and Pechenik 1997; Jarrett 2003; Thiyagarajan et al. 2005; Emlet and Sadro 2006). Previous research suggests that environmental factors, such as chlorophyll-*a* concentrations (a proxy of food availability for larvae), may drive larval quality variations and, consequently, recruitment dynamics (Barbosa et al. 2016), thereby influencing the establishment of marine invertebrate populations (Olivier et al. 1996; Hunt and Scheibling 1997; Jennewein and Gosselin 2013). Our findings provide evidence for these fitness advantages, as settlement on preferred rugose habitats increased survival by up to 40% and enhanced growth rates by approximately 20 µm day^-1^. Such structural heterogeneity likely functions as a fitness refuge, providing shelter from predation and buffering abiotic stressors during emersion, such as thermal fluctuations and solar radiation (Jenewein and Gosselin 2013).

Recruitment patterns during the second 16-day period were associated with high-quality larvae, suggesting increased settlement competency. This shift aligns with the general absence of low-quality larvae (index <1.7), indicating that ‘desperate larvae’ were not present during this phase. However, despite the presence of high-quality larvae in Period 2, substrate selectivity was no longer observed. This loss of discriminatory behavior may be related to the progressive development of biofilm on the settlement plates after 10 days. Biofilm formation follows a successional pattern, beginning with bacterial microfouling and evolving into an organic matrix that supports a diverse community, including microalgae (mainly diatoms), sponges and ciliate protists (Kristensen et al. 2008). While microbial biofilms often facilitate settlement by enhancing adhesion (Zardus et al. 2008), their effects are context-dependent and vary based on biofilm composition (Qian et al. 2007). Differences in bacterial community composition within biofilms have been shown to alter or even inhibit barnacle settlement (Lau et al. 2005), which likely explains the absence of habitat selectivity observed in high-quality cyprids during this second period.

These findings underscore the critical role of larval quality in shaping recruitment success for habitat-forming benthic invertebrates. The observed variation in settlement selectivity highlights the interplay between individual physiological state (energetic reserves) and environmental factors (substrate condition) in modulating settlement dynamics. Therefore, recruitment models should integrate larval quality together with the spatial and temporal heterogeneity of environmental conditions to improve predictions of settlement success and population connectivity (Pechenik et al. 1998; Cowen et al. 2000; Levin 2006). Future research should further explore the interactions between larval quality, biofilm composition, and settlement success across varying environmental conditions. Investigating how environmental stressors, such as temperature and food availability, influence larval lipid reserves could provide critical insights into the mechanisms driving settlement variability. Integrating larval quality assessments into ecological frameworks can refine recruitment predictions and strengthen conservation strategies for benthic marine populations.

